# Ranking Antibody Binding Epitopes and Proteins Across Samples from Whole Proteome Tiled Linear Peptides

**DOI:** 10.1101/2023.04.23.536620

**Authors:** Sean J. McIlwain, Anna Hoefges, Amy K. Erbe, Paul M. Sondel, Irene M. Ong

## Abstract

Ultradense peptide binding arrays that can probe millions of linear peptides comprising the entire proteomes or immunomes of human or mouse, or numerous microbes, are powerful tools for studying the abundance of different antibody repertoire in serum samples to understand adaptive immune responses. There are few statistical analysis tools for exploring high-dimensional, significant and reproducible antibody targets for ultradense peptide binding arrays at the linear peptide, epitope (grouping of adjacent peptides), and protein level across multiple samples/subjects (I.e. epitope spread or immunogenic regions within each protein) for understanding the heterogeneity of immune responses. We developed HERON (**H**ierarchical antibody binding **E**pitopes and p**RO**teins from li**N**ear peptides), an R package, which allows users to identify immunogenic epitopes using meta-analyses and spatial clustering techniques to explore antibody targets at various resolution and confidence levels, that can be found consistently across a specified number of samples through the entire proteome to study antibody responses for diagnostics or treatment. Our approach estimates significance values at the linear peptide (probe), epitope, and protein level to identify top candidates for validation. We test the performance of predictions on all three levels using correlation between technical replicates and comparison of epitope calls on 2 datasets, which shows HERON’s competitiveness in estimating false discovery rates and finding general and sample-level regions of interest for antibody binding. The code is available as an R package downloadable from http://github.com/Ong-Research/HERON.

## 1. Introduction

The technology for high-dimensional identification of specific antibody binding repertoire has significantly improved over the last decade, allowing for the determination of antibody binding to ∼6 million peptides simultaneously with peptide array technology to probe every mouse or human protein using 16-mer peptides with 1, 2 or 4 amino acid (a.a.) tiling to identify antibody targets from serum or plasma [1]. This extremely high-dimensional, high-throughput method for antigen-specific immune profiling can enable detection of immune responses to infection or vaccines, and potentially study the precise targeting of tumors by one’s own immune system with or without immunotherapy/combination therapy. There are many existing methods for analyzing microarray and peptide array data [2-15], including pepStat, which was designed to analyze different viral strains for a single protein, and a few methods for analyzing these ultra-dense, high-dimensional array data [13, 15, 16], but additional methods are necessary to advance understanding of immune response as antibody binding signals from arrays represent an aggregate of complex biological and biochemical interactions. Antibodies may bind to peptides via various mechanisms of interaction based on the antibody’s antigen binding site and amino acid configuration of the peptides, affecting binding affinity, and each protein may have multiple epitopes that are bound by antibodies, amplifying the immune response to the protein. These considerations are important when studying immune responses involved in diseases, including infectious, auto-immune, or neoplastic. To the best of our knowledge, there are currently no existing methods that can simultaneously identify and rank antibody binding responses at different scales, i.e., to linear peptides, epitopes (defined here as antibody bound region of adjacent linear peptides or probes tiled within a protein) and proteins, for diagnostics or treatment; or for understanding the heterogeneity of immune response within an individual or across a population.

Existing methods for analyzing antibody binding of peptide array data such as pepStat [2], estimates the false discovery rate (FDR) at a peptide-level and reports subject-level statistics. One Bayesian model, pepBayes, has been shown to be potentially superior to pepStat, however, it was also designed to analyze a single protein from multiple related viral strains and does not take into account the adjacent probes across the protein [3]. Another Bayesian approach does include provision for handling sequential probe signals by using a latent autoregressive component, however the authors propose a limit in the problem size (300 peptides and 50 samples) when using their implementation [17]. MixTwice [13] is a statistical method that utilizes additional power to detect significant probes by using local FDR. However, none of these approaches ensure a reproducible way to rank epitopes and score both the epitopes and proteins when analyzing whole proteome peptide arrays with ∼6 million unique sequence probes.

We developed an algorithm, HERON (**H**ierarchical antibody binding **E**pitopes and p**RO**teins from li**N**ear peptides), which is an R [18] package available on GitHub, for analyzing ultra-dense peptide arrays to identify and rank significantly bound linear peptides, epitopes, and proteins. The overall workflow is shown in **Figure 1**. Our approach builds on existing approaches [2] including clustering methods to locate contiguous probes (i.e., epitopes) with high binding affinity, and meta-analysis methods from Fisher and others [19-25] to: 1) allow for reliability and reproducibility with granularity in confidence level for making antibody binding calls at different thresholds, 2) ensure that we identify probes that are more highly bound in post-treatment (experimental) samples compared to pre-treatment samples (control), and that we give more weight to those with higher overall binding, 3) identify consecutive overlapping probes with high binding signal and categorize the shared amino acid (a.a.) sequences represented by those highly recognized probes as epitopes based on specified thresholds, and 4) identify proteins due to epitope spread (i.e., identify proteins recognized by distinct epitope binding in different regions of the protein by sera from multiple samples, not necessarily at the same place on the protein).

**Figure 1.**
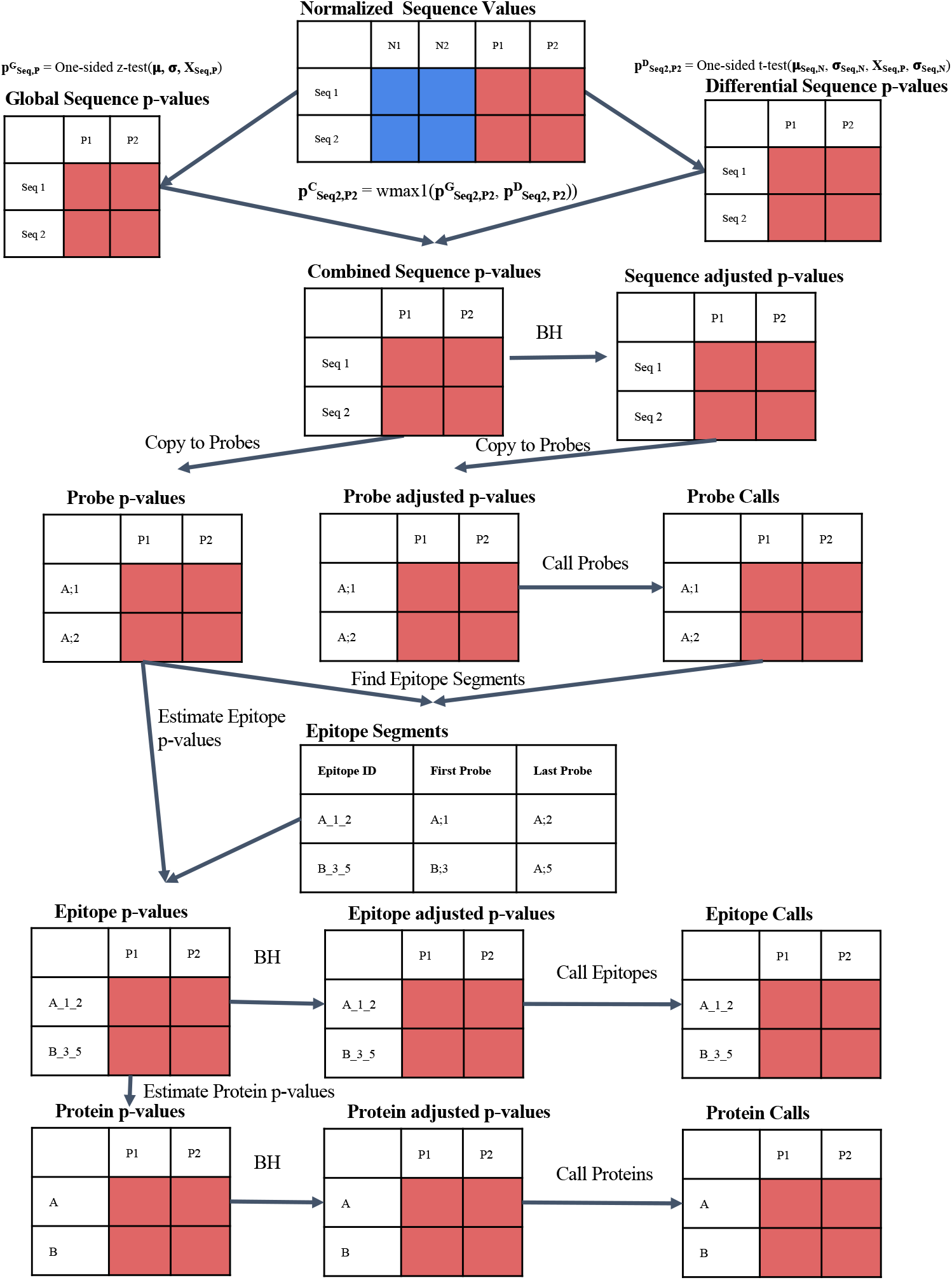
HERON workflow for processing peptide binding array data. Workflow illustrating algorithm that takes as input a normalized matrix of sequence probes and samples and outputs probe-level, epitope-level, and protein-level differential antibody binding calls on each sample. The combined sequence p-value are calculated by combining a global p-value from a one-sided z-test with a differential p-value from a one-sided t-test using the Wilkinson’s max (wmax1). The N1 and N2 columns (dark-blue colored) indicate the pre-treatment sample values and output and the P1 and P2 columns (dark-red colored) indicate the post-treatment sample values and output. After adjustment of the sequence p-values using Benjamini-Hochberg, the raw sequence p-values and adjusted sequence p-value are copied to the respective protein probe locations. An FDR cutoff is used to identify the probes significantly differentially bound by antibodies. The epitope regions, where the Epitope ID is defined by Protein_FirstProbe_LastProbe of the epitope, are detected using a segmentation algorithm, and the found segments are scored using meta-analyses methods to combine to contained probe-level p-values, corrected using BH, and an FDR threshold is used to identify significant differentially bound antibodies. Finally, the protein scores are determined from the associated protein’s epitope p-value(s) using meta-analyses methods, corrected using BH, and then called using an FDR threshold.

To ensure that probes that are more highly bound in post-treatment (experimental or positive) compared to pre-treatment (control or negative) are identified, and that probes with higher overall binding are given more weight or a smaller estimated p-value, our algorithm calculates probe-level p-value scores for each post-treatment sample by combining t-test scores for differential expression (one sided test above control) and global z-tests normalized for relative height compared to all of the data in the experiment (**Figure 1**).

In order to identify common and unique epitopes across individual samples we find regions of consecutive probes across the tiling of the proteins that are consistent with their calls across positive samples by making initial antibody binding calls on the probe-level adjusted p-values based on a threshold, then utilizes one of three approaches for identifying epitopes, which are consecutive probes tiled across a single protein. After finding the epitopes, we then calculate a score for all the epitopes using a respective meta p-value estimation method [19-25] and adjust for false discoveries using Benjamini-Hochberg (BH) [26]. A similar approach using meta-analysis is applied to epitope level calls at the protein level to calculate scores for all the proteins.

In this report, we measured the performance of our method on two datasets to illustrate its utility. The goal of the first dataset (COVID-19) was to identify diagnostic epitopes across the landscape of the SARS-CoV-2 viral proteome as detected in human serum samples from individuals following SARS-CoV-2 infection [11]. The goal of the second dataset (Melanoma) was to study the landscape of antibody responses to the whole mouse proteome of genetically identical mice before cancer and after curative immunotherapeutic *in situ* vaccine treatment [27] and rechallenge with a related tumor and to rank the responses for study and validation. For evaluation of the COVID-19 dataset, we show performance measures for finding significant epitopes by comparing them with the regions identified using a t-test method from the SARS-CoV-2 manuscript [11] and for the Melanoma dataset, we measured correlation between technical replicates using different parameter settings.

## 2. Materials and Methods

### 2.1 Datasets

The COVID-19 dataset [11] compares analyses of serum samples from individuals following proven infection with SARS-CoV-2 (COVID+) with serum samples from uninfected individuals (COVID-); this dataset was downloaded following the instructions from https://github.com/Ong-Research/UW_Adult_Covid-19. There are 60 (20 COVID-, 40 COVID+) samples in total, with 118,651 unique sequence probes mapped to 470,086 16-mer probes for 387 proteins with a tiling of 1 amino acid (a.a.) and up to 5 replicates on the array. The data were preprocessed using pepMeld [28] and the resulting sequence matrix was quantile normalized at the sequence probe level.

The Melanoma dataset compares naïve mouse serum to mice that have successfully rejected a tumor after immunotherapy and also showed immune memory when presented with a rechallenge of the same or a similar tumor type. These mice had an established B78 melanoma tumor, have been treated with a potent immunotherapy treatment consisting of 12 Gy external beam radiation followed 5 days later by 5 consecutive days of intratumor injections of an immunocytokine which targets the GD2-expressing tumor via a monoclonal antibody and delivers IL-2 directly to the tumor site to elicit a strong and long-lasting anti-tumor immune response.

The Melanoma dataset was collected by Hoefges et al. 2023 [29] and can be found in the Immport repository under accession number SDY2162 (https://www.immport.org/home). The dataset consists of 11 biological (serum) samples tested for recognition of 6,090,593 16-mer unique sequence probes mapped to 8,459,970 protein probes using a mixed tiling of either 2 a.a. or 4 a.a. with a total of 53,640 individual proteins. Of the 11 samples, 5 samples were from naïve mice, (referred to as “pre-treatment”; i.e., prior to tumor introduction or immunotherapy), and 6 were immune samples (referred to as “post-treatment”; i.e., they were tumor-bearing mice that underwent radiation and immunotherapy treatment to become tumor-free. Following tumor-clearance, the mice were rechallenged with the tumor line to test immune memory). Two of the immune samples had technical replicates: one (sample B2) had replicates of the same Immune serum sample tested on the same array (performed within 24 h of each other) while the other (sample PD1) had the replicates of the same immune serum sample on identical arrays but performed ∼ 1 year apart from each other. In total, there were 13 serum samples, 5 from naïve mice, 8 from 6 immune mice, (including the 2 replicates for B2 and PD1). The data were pre-processed by Nimble Therapeutics, quantile normalized, and then smoothed using a sliding average mean window across the protein location of +/-8 a.a.

### 2.2 Algorithm

The R package, HERON, can be downloaded and installed from GitHub (http://github.com/Ong-Research/HERON). The overall workflow of the algorithm is illustrated in Figure 1 including the underlying approach for finding significant probes (**Figure 1B**). Herein, we describe the methods for finding consecutive blocks of probes on the protein, which we call epitopes, as they can be used to identify the exact sequence of 2-16 consecutive amino acids, shared by consecutive probes, all recognized by the same antibody, and thereby corresponding to an epitope recognized by an antibody [29]. Given the found epitopes, we then describe how to estimate the significant epitopes and corresponding proteins.

#### 2.2.1 Estimating probe-level p-values for each positive sample

HERON was designed to analyze ultradense peptide binding arrays with multiple protein sequences, e.g., whole mouse or human proteomes, and the goal is to rank or calculate a p-value/false discovery rate (FDR) for each linear peptide probe for each positive sample and summarize ranking at different resolutions (at epitope and protein levels) for multiple positive samples. As shown in **Figure 1B**, we first calculate a global p-value using a distribution function for all of the data provided in the experiment, then a differential p-value using a one-sided t-test for each positive sample is calculated assuming that the standard deviation is the same between groups using the pre-treatment or post-treatment samples. Parameters for generating probe-level p-values can be adjusted and one can use either or both of the global and differential tests when calculating significance for peptides to identify probes that are more highly bound in post-treatment (experimental or positive) compared to pre-treatment samples, or give more weight to those with higher overall binding, respectively. Our approach calculates a combined p-value from the global p-value and differential p-value using Wilkinson’s max meta p-value method [20, 24, 30, 31]. **Supplementary Figure 1** shows the estimated global p-values (**S1A**), differential p-values (**S1B**), combined global and differential p-values (**S1C**), and adjusted p-values (Benjamini-Hochberg, **S1D**) versus the peptide array signal after applying the HERON workflow on data for a positive sample from the Melanoma dataset, where a low p-value/FDR indicates an increased probability that both the global and differential hypotheses are rejected.

#### 2.2.2 K of N Calls and One-Hit Filter

The linear probe-level p-values are then adjusted for false discovery rates using the Benjamini-Hochberg algorithm. Next the significance values for the linear peptides are calculated and an FDR threshold cutoff is used to identify the set of antibody-bound probes (which we will also refer to as “calls” or “called” probes) for each positive sample. HERON also reports the number of samples that have an adjusted p-value less than the specified threshold, i.e., reporting a number, K samples that indicated antibody binding for the called probes, out of a total of N samples. To filter out inconsistent calls due to spurious noise (or non-specific signals), the algorithm has a one-hit filter option for removing probes that are called as bound by antibodies that do not have a supporting consecutive probe call in the same serum sample and does not have a call for the same probe in another serum sample. The one-hit filtering is a soft procedure, where the raw and adjusted p-values are set to 1 for the filtered one-hit probe. These procedures are also applied to epitope- and protein-level probes after the steps described below.

#### 2.2.3 Epitope Finding

Once the set of antibody-bound probes are identified, epitopes can then be found using three different segmentation algorithms: unique, hierarchical, or skater.

The unique method iterates through all the samples, finding consecutive runs of probes that were called on the same protein, then combines all the regions found across the samples into a single list, and reports the unique set of blocks/epitopes. The method permits overlapping blocks. **Figure 2A** illustrates the “unique” algorithm on an example set of called probes.

**Figure 2.**
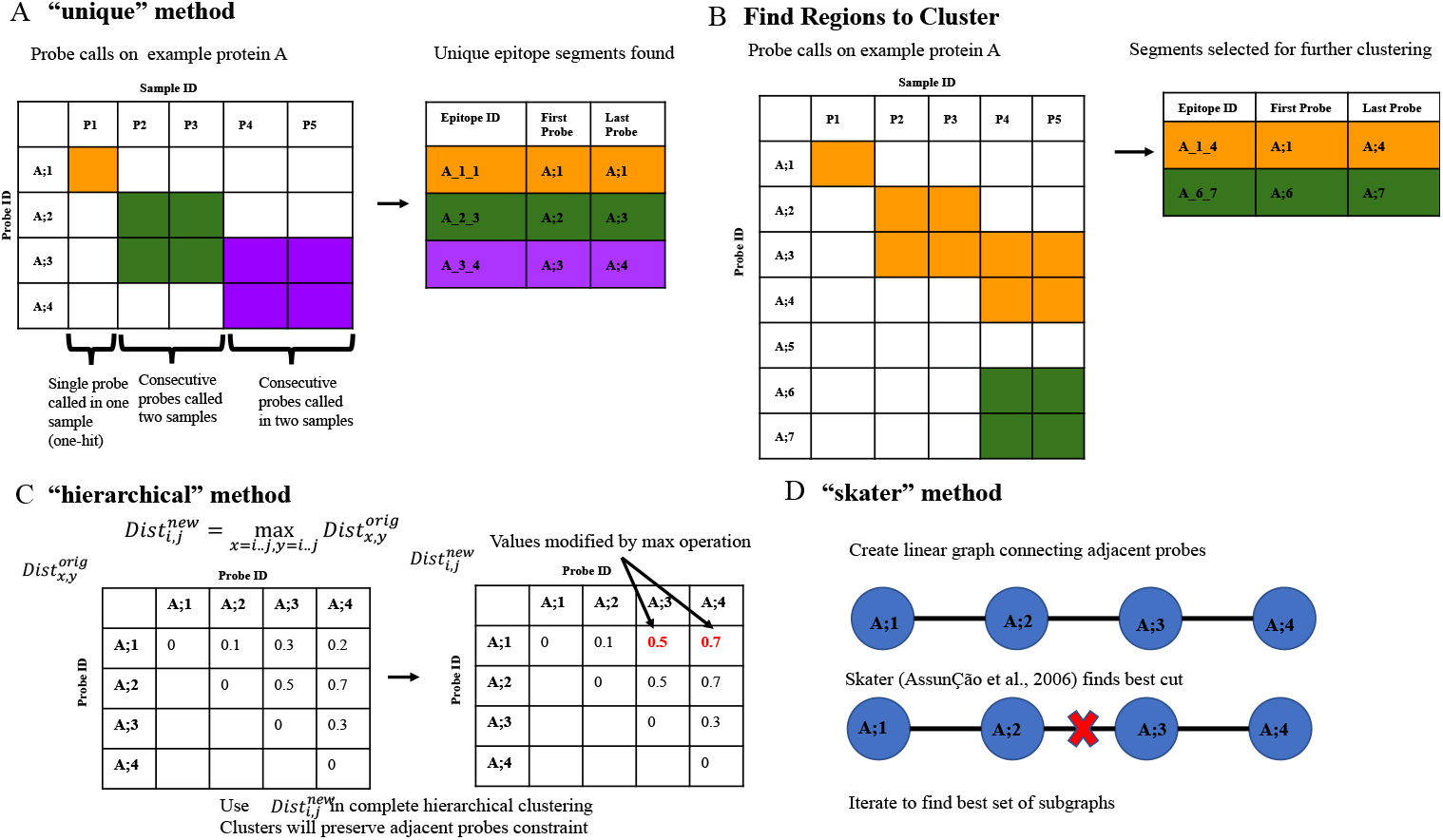
Illustration and Example of epitope finding algorithms implemented in HERON. (A) Illustration of the “unique” method, which finds the unique set of grouped probes called from each post-treatment sample. The cells highlighted in orange, green, or purple are the called probes. The table to the right of the illustration is filled in with the color corresponding to the outlined boxes in the illustration (orange, green and purple) to indicate the called epitope probe regions in the illustration. Epitope ID is defined by Protein_FirstProbe_LastProbe of the epitope. (B) Illustration of how regions are found to further cluster using either the hierarchical or skater method. The orange and green highlighted boxes mark the probes that are adjacent and the table on the right indicates the larger groups of probes, in the corresponding orange and green, that will be further segmented using the clustering methods. (C) Illustration of the hierarchical clustering method, which adapts hierarchical clustering to find consecutive probes with consistent call patterns across samples. The matrix on the left is the original distance matrix calculated from the dissimilarity of the probe calls or scores. The matrix on the right is the new matrix after applying the max operation. The two red numbers indicate the cells that were changed due to the max operation. (D) Illustration of the “skater” method which decomposes a linear graph of consecutive probes to find consistent call patterns across samples.

Both the hierarchical clustering (hclust call in base R) [32] and skater (spdep R package) methods [33, 34] in HERON first finds regions within each protein where consecutive probes are called in any positive sample (**Figure 2B**). These groups of called probes are then segmented using either the hierarchical (**Figure 2C**) or skater (**Figure 2D**) clustering method and the average silhouette score is used to determine the optimal number of clusters or graph cuts respectively.

To calculate similarity or dissimilarity between probes in relation to their significance scores or calls for the post samples, HERON provides the option to utilize a “binary” or a “z-score” method. The binary method uses the probe calls as labels (true/false for bound/unbound by an individual serum sample) for which a hamming distance can be calculated between the probes. The z-score method converts the probe p-values to a one-sided z-score before clustering using the Euclidean distance. After building a distance matrix for all possible pairs of probes on the protein, we then use a clustering algorithm to find groups of consecutive probes on the protein that are similar in their calls or significance values across the post-treatment samples. For both the hierarchical and skater segmentation implementation we use the maximum average silhouette score to determine the number of clusters (hierarchical clustering) or cuts (skater). Ties in the silhouette score are broken by selecting the clustering with the maximum number of clusters or cuts. Other different distance metrics can also be used for measuring the distance between elements; however, our algorithm used the Euclidean distance with the z-score clustering and an average hamming distance for segmenting using the binary calls.

Using the hierarchical clustering algorithm from hclust, HERON first calculates the distance matrix as usual. For each identified cluster, a distance matrix is derived by setting each i/j element to the maximum pairwise distance between probes starting at protein position i and ending at protein position j for the group of probes to be segmented. Complete hierarchical clustering is then performed using the distance matrix to find clusters that are contiguous probes within the protein (**Figure 2C**).

The skater algorithm is used to find consecutive probe regions with similar signal patterns across the samples. Since skater allows graph constraints on the clustering process, HERON defines a linear graph where an edge is introduced between each peptide that is the adjacent peptide within the protein tiling. Each iteration of the skater algorithm finds the best cut in the graph using the calculated distance metric between the probes. After iterating through all possible cuts to find the cut that gives the best average silhouette score, the epitope regions are then defined by the remaining connected nodes in the graph (**Figure 2D**).

#### 2.2.4 Epitope and Protein p-values

An epitope level p-value is calculated by using a meta p-value method with each positive sample’s linear probe p-values for the probes that are contained in that epitope. Similarly, the protein level p-value is determined using a meta p-value method based on the epitope p-values found across the protein for each positive sample. Many meta p-value estimation methods exist [19-24, 35, 36], including fisher, and each has its own characteristics depending upon the type of underlying hypothesis to be tested.

For epitopes, we chose a meta p-value method that requires most, if not all, of the peptide probes within the epitope block to have significant values, such as Wilkinson’s max (wmax) [20, 24, 30, 37]. To relax the requirement that all probe p-values within an epitope be significant, Wilkinson’s max can be calculated on the n^th^ maximum p-value (I.e., wmax2 for the 2^nd^ highest p-value within the epitope). In cases where Wilkinson’s max is used and the number of probes in the epitope is fewer than n peptides, HERON conservatively assumes that the p-value was 1. Due to the inherent possibility of dependency between the signals of adjacent probes, HERON also allows the use of the harmonic mean (hmp) [19] and the Cauchy (cct) [23] meta p-value methods, which have been shown to be tolerant to inter-dependencies between the elements for which the p-values are combined.

At the protein level, our goal was to identify proteins where at least 1 (or more) epitopes are significant, a meta p-value method of choice would be the minimum + Bonferroni correction (min_bonf) or the Wilkinson’s minimum (wmin) or Tippett’s method [20, 24, 30, 37]. Using the nth minimum (I.e., wmin2 for the 2^nd^ smallest epitope p-value within the protein) would be a more stringent requirement, where at least n epitopes have to be significant. In the case where there are fewer than n epitopes with p-values, HERON tries to call Wilkinson’s min with the (n-1)^th^ p-value iteratively. If there is only one p-value, then Wilkinson’s n^th^ min will just return the single p-value.

## 3. Results

The performance of the algorithm and optimal parameter settings were tested on two different datasets, COVID-19 and Melanoma. Performance was evaluated by testing the correlation between technical replicates for repeatability of calls, finding peptides that ensure reproducibility with a validation assay such as ELISA, and epitope boundary finding parameters as described in the next sections.

### 3.1 Comparison of HERON performance on COVID-19 Dataset

We compared HERON’s performance on a COVID-19 dataset, using parameters that were similar to the t-test workflow used to identify diagnostic peptides [11]. We utilized the following significance parameters: an absolute shift on the differential t-test of one, where one would mean the difference between the current COVID+ sample’s normalized fluorescent value from the Nimble system sample and the average normalized values from the COVID-samples is significantly different by more than 2-fold, an adjusted p-value threshold <0.01, one-hit filter, hierarchical clustering with binary and hamming distance scores, Williamson’s max and Tippett’s for epitope and protein meta p-values estimation respectively. Probe, Epitope, and Protein calls were made if 25% of the COVID+ samples were called at the adjusted p-value threshold of <0.01. We also use the differential t-test (not the global z-test) to resemble the processing done in the Heffron et al. 2021 paper. Also, to compare against other methods, we also ran pepStat using the method’s normalization method and no smoothing and an FDR cutoff 0.01.

Each segmentation result and subsequent calls at the probe, epitope and protein levels are compared against the previously reported regions [11] (**Figure 3A**). For the epitope-level, we convert the selected epitopes back to the list of probes that are contained within each epitope, and then compare the probes between the two methods. The overlaps and probes unique to the Heffron et al. 2021 and the HERON method are provided in **Supplementary Table 1**. All but 7 probes previously reported were identified by HERON indicating good agreement between the probe calls. The differences between the probes called can be due to 1) relaxation for every COVID+ sample that gets called separately using the standard deviation from the COVID-samples, 2) the Heffron paper uses a strict 2-fold change cutoff whereas HERON is using a less stringent method that incorporates the 2-fold change as part of the differential p-value calculation. Looking at the probes, we find that of the 7 missed by HERON, 4 are on the edge of achieving the 25% cutoff (9 out of 40 COVID+ samples). Also, all 7 of the probes missed by HERON are part of short (1-2 probes) epitopes. Conversely, of the 145 probes identified in HERON but not in Heffron et al., 56 are on the edge of not being called (exactly 10 out of 40 COVID+ samples).

**Figure 3.**
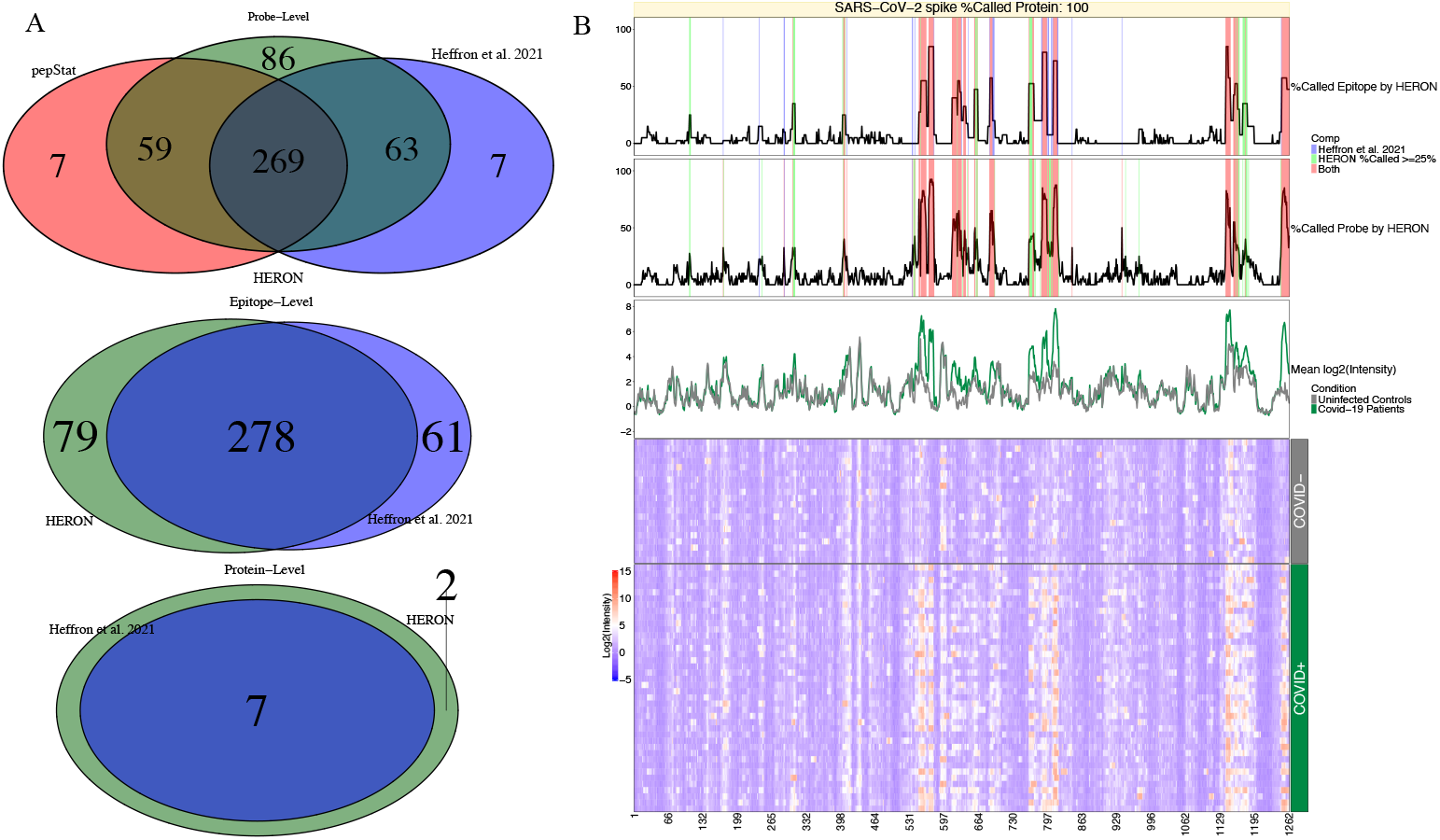
Venn Diagram comparisons for COVID-19 dataset and Heatmap of SARS CoV-2 spike protein. (A) Venn diagrams of the linear peptide probe-level (top), epitope-level (middle), and protein-level (bottom) where the green circle represents the calls made by HERON, the red circle represents the probe calls made by pepStat [2], and the blue circle represents the calls made by Heffron et al. 2021 [11]. (B) Annotated heatmap [38, 39] of SARS CoV-2 spike protein. The heatmap (at the bottom) depicts the normalized intensity values for the probes (x-axis) tiled across the SARS-CoV-2 membrane protein and the individual patient serum samples (y-axis) with red representing high antibody (Ab) binding and blue representing low Ab binding. The top line plot indicates the percent of COVID-19 positive (COVID+) samples that were called as significantly differentially bound at least 2-fold over the average of the COVID-19 negative (COVID-) samples and after calculating the meta p-values on the epitope-level, the middle line plot indicates the percent of COVID+ samples called significantly 2-fold over the COVID-samples on the probe-level, and the bottom line plot shows the average signal between control (COVID-) and positive samples. In these line plots, the % Called Probe by HERON and % Called Epitope by HERON are highlighted in red if both the HERON and Heffron et al. 2021 method called the same probes or group of probes, blue if only the Heffron et al. 2021 method called the probe or epitope, and light green if the probe or epitope was called by only the HERON method.

With the exception of 7 probes, most of the probes called by pepStat were also called with HERON. The six of seven probes found by pepStat appear to be on the consecutive edges of probes that would have been called in an epitope block and one probe in orf1ab that appears to be novel.

Figure 3B is a heatmap of the spike protein with line graphs showing the probe and epitope annotations of the SARS-CoV-2 spike protein from the heat map displayed across the top. The epitope and probe annotations also display the agreements and disagreements between HERON and the epitopes and probes as previously reported [11]. The differences between the epitope calls are most likely due to the method for finding epitope segments and the meta p-value method used to score them. The overlaps and epitope probes unique to the Heffron or HERON methods are provided in Supplementary Table 2. Of the 61 epitope probes that were missed by HERON, 43 are due to an epitope extension, where HERON finds a longer epitope than the previous method, but results in a loss of significance. 3 of the 43 extensions are on the edge of 25% (9 out of 40 COVID+ samples). The remaining 18 have matching epitopes in HERON but are shifted or fragmented and also have a loss of significance. Subsequently, of the 79 epitope probes found in HERON but not in the previous method, 49 are extensions (with 9 on the edge, i.e., 10 out of 40 COVID+ samples called), and 21 are from new epitopes (with 6 of the 21 on the edge of being called. These results indicate that HERON can find epitopes that can be more specific to a subset of subjects, which warrants additional study to further improve the epitope segmentation process.

The protein level Venn diagram at the bottom in Figure 3A shows HERON selected additional proteins than the previous method [11]. Since the requirement on the protein-level is to have one or more significant epitopes and many potential epitopes are detected on the proteins, the proteins selected in the COVID dataset will be permissive in the number of proteins called.

### 3.2 Repeatability of Probe, Epitope, and Protein Significance Values with the Melanoma Dataset

We studied the repeatability of the p-value on the probe, epitope, and protein level by looking at the correlation between the significance values obtained on technical replicate samples from the murine Melanoma immunotherapy dataset. The Pearson correlation of the -log_10_(FDR) between two technical replicates: one single immune serum sample was divided into two identical aliquots which were tested in parallel with parallel data sets collected in the same array assay (B2), or a separate single immune serum sample divided into two identical aliquots which were tested on similar arrays, independently with parallel data sets collected in two separate similar arrays that were performed approximately one year apart (PD1).

For the parameters specific to probe p-value calculations, we chose three different levels of significance on the global z-test p-value coupled with the FDR cutoff used (Inclusive – global with a standard deviation (sd) shift of 3 and adjusted p-value cutoff of <0.2, Moderate – global sd shift of 6 and adjusted p-value cutoff of <0.05, and Restrictive – global sd shift of 10 and adjusted p-value cutoff of <0.01) and investigated with or without the use of the one-hit filter. For the epitope p-values parameters, we chose five different ways of finding epitopes (unique, hierarchical clustering with binary calls and hamming distance (hbh) or z-score with Euclidean distance (hze), skater with binary calls and hamming distance (sbh), and skater with z-score and Euclidean distance (sbz). The epitope meta p-value was either Wilkinson’s max (wmax1), Wilkinson’s 2nd max (wmax2), Fisher (fisher), harmonic mean (hmp), or Cauchy (cct). For protein meta p-values, we choose either Fisher (fisher), Tippetts/Wilkinsons min (wmin1), Wilkinson 2nd min (wmin2), or min with a Bonferroni correction (min_bonf). Different adjusted p-value cutoffs could be used for the probe, epitope, and protein levels. Our results have tied these three parameters to the same value.

We calculate the Pearson correlation for the between the technical replicates of B2 and PD1 using the -log10(adjusted p-values) for the probes, epitopes, and proteins using an exhaustive search of the parameters mentioned above. The correlation results are presented in **Supplementary Table 3**.

Looking at the average correlations between the probe-level p-values of the technical replicates of B2 and PD1 (**Supplementary Table 4**) and marginalizing across the one-hit filter results, we found that using more stringent statistical filtering improves the technical replicate Pearson intercorrelation of the probe –log_10_(adjusted p-values) (Inclusive – 0.806, Moderate – 0.841, and Restrictive – 0.846). By marginalizing across the statistical parameters, we also found that using the one-hit filter slightly improves the average replicate intercorrelation (Without – 0.824, With – 0.837). To find a good setting with good repeatability across the probe level significance values used, we then averaged the average correlation for the probe, epitope, and protein across the inclusive, moderate, and restrictive statistical parameters. The overall correlation results are reported in **Supplementary Table 5** and the top 10 parameter settings are summarized in **Figure 4A**. Looking at the top 10, it appears that the Wilkinson’s max on the 2^nd^ highest p-value for epitopes and either Tippett’s or min+Bonferroni for protein meta p-values using the skater or hierarchical clustering segmentation methods on the binary calls achieve the highest average correlation across the significance levels and the two technical replicates.

**Figure 4.**
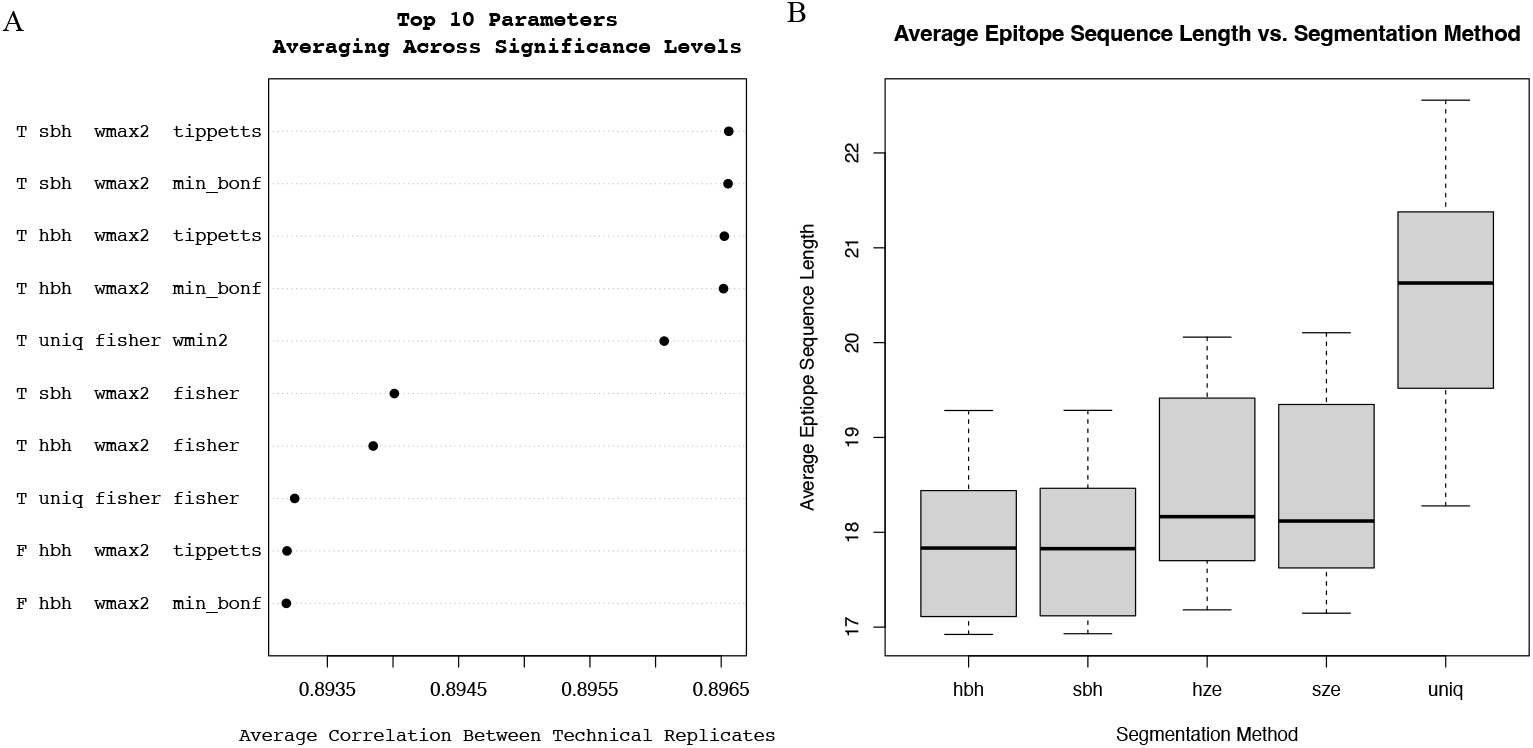
Performance Results for Melanoma Dataset. **(A)** Average Correlation between technical replicates for the Top 10 algorithm parameter settings. Each row-label is as follows: One hit filter (T - True or F – False indicates whether filter is used or not), Segmentation method (*uniq* – unique set of epitopes found across all post samples, *hbh* - hierarchical clustering with binary calls with hamming distance, *sbh* - skater with binary calls and hamming distance, and *sbz* - skater with z-score and Euclidean distance), epitope meta p-value method (*wmax2* – Wilkinson’s max on the 2^nd^ largest p-value, *fisher* – Fisher’s method), protein meta p-value method (*tippetts* – Tippett’s or Wilkinsons’s min on the 1^st^ smallest p-value, *min_bonf* – finds the minimum p-value then correct by the number of epitopes on the proteins using Bonferroni), *wmin2* – Wilkinson min on the 2^nd^ smallest p-value). (B) Average Epitope Sequence Length vs. Segmentation Method. Box of the average sequence length that results from stitching the contained probe sequences for the detected epitope. For each segmentation method, the box is calculated using all of the average epitope sequence lengths for the remaining parameters, keep the epitopes that have at least one sample called.

The choice of segmentation method can also determine the length of the epitope (Table 4B). It appears, on average, that the “unique” (uniq) segmentation method gives longer epitopes, while the skater or hclust segmentation methods using the binary calls obtains shorter epitopes. Looking at Figure 4A, the shorter epitopes maybe indeed be more accurate since those methods are within the top 2 parameters for highest average technical correlation.

**Supplementary Figure 2** shows technical replicates as scatterplots for each of the probe, epitope, and proteins for the best overall average correlation using the Moderate global standard deviation shift and FDR cutoff values.

### 3.3 Probe, Epitope, and Protein Counts with Different Significant levels on the Melanoma Dataset

We used the best correlation parameters to make calls on the Melanoma dataset for which the technical replicates were averaged together at the inclusive, moderate, or restrictive significance levels to obtain calls at the probe, epitope, and protein level. The significance calls for each sample and level for the moderate significance values are displayed as a complex upset plot [40, 41] in **Figures 5A – Probe-level, 5B – Epitope-level, and 5C - Protein-level**. The number of probes, epitopes, and proteins recognized by 4, 5 or 6 of the 6 immune mice was substantially lower than the number of epitopes mutually recognized by multiple individuals. **Figure 6** presents data for the LEM containing protein 3 (Lemd3), illustrating where two epitopes were found in two different positive samples (1 out of 6, ∼16%), which resulted in a K of N of 2 (33%) at the protein level. **Supplementary Figure 3** depicts the Hemicentin-1 (Hmcn1) protein, which had 11 epitopes called in different regions with different numbers of positive samples called.

**Figure 5.**
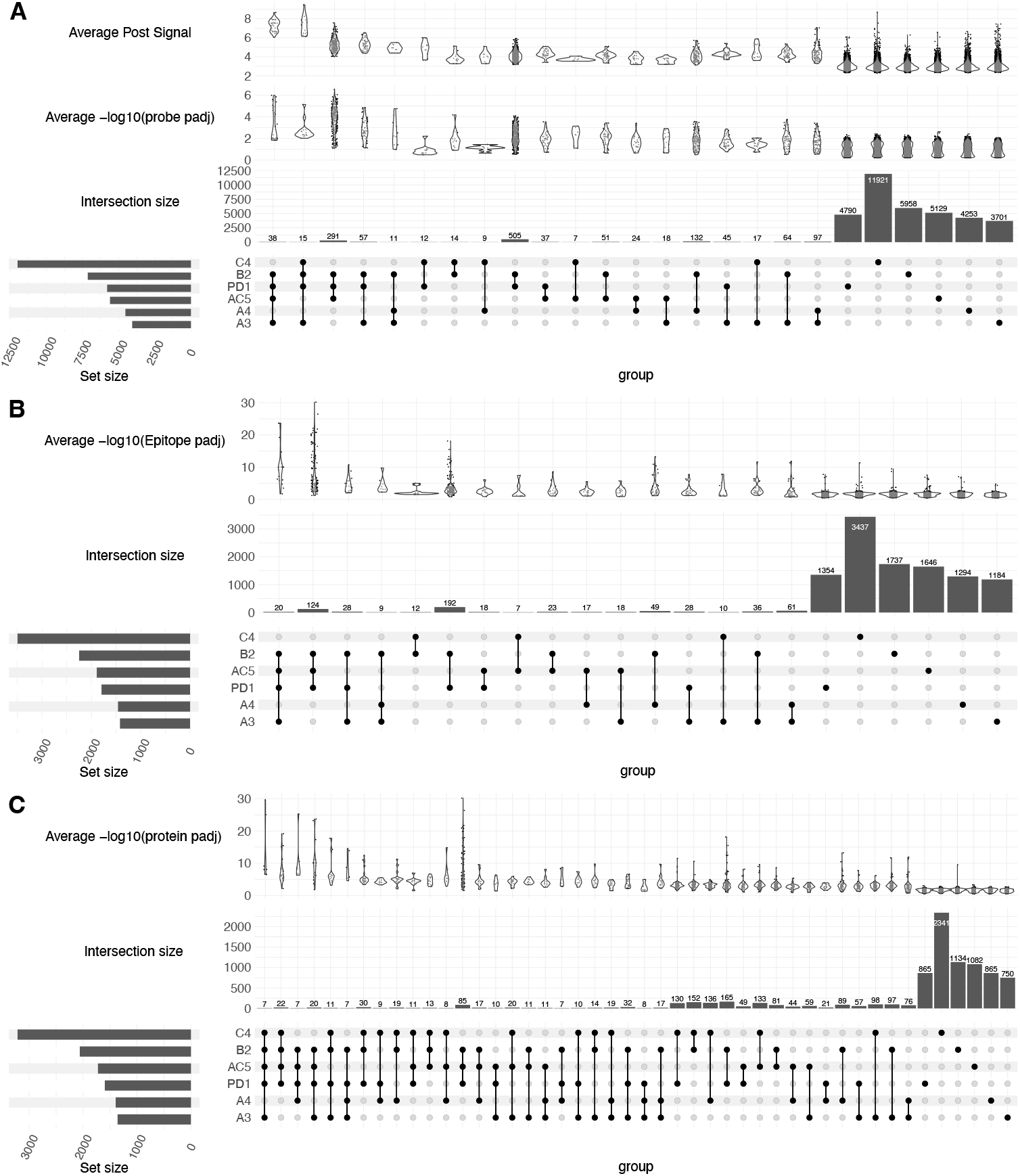
Upset Plots with Average Post Signal and -log10 adjusted p-values. Upset plot of the probes (A), epitopes (B), and proteins (C) called for each sample using the moderate significance level (min set size = X, sorted by the decreasing number of categories in the intersection set). AC5, A3, C4, A4, B2 and PD1 are the designations for the individual mice cured of melanoma, that provided the 6 separate immune serum samples tested here. For the probe plot (A), the top row of violin plots shows the average post signal for each intersection, the middle row of violin plots shows the average of -log_10_(probe adjusted p-values) for each intersection. The epitope (B) and protein (C) violin plots are the respective -log_10_(epitope adjusted p-value) and -log10(protein adjusted p-values).

**Figure 6.**
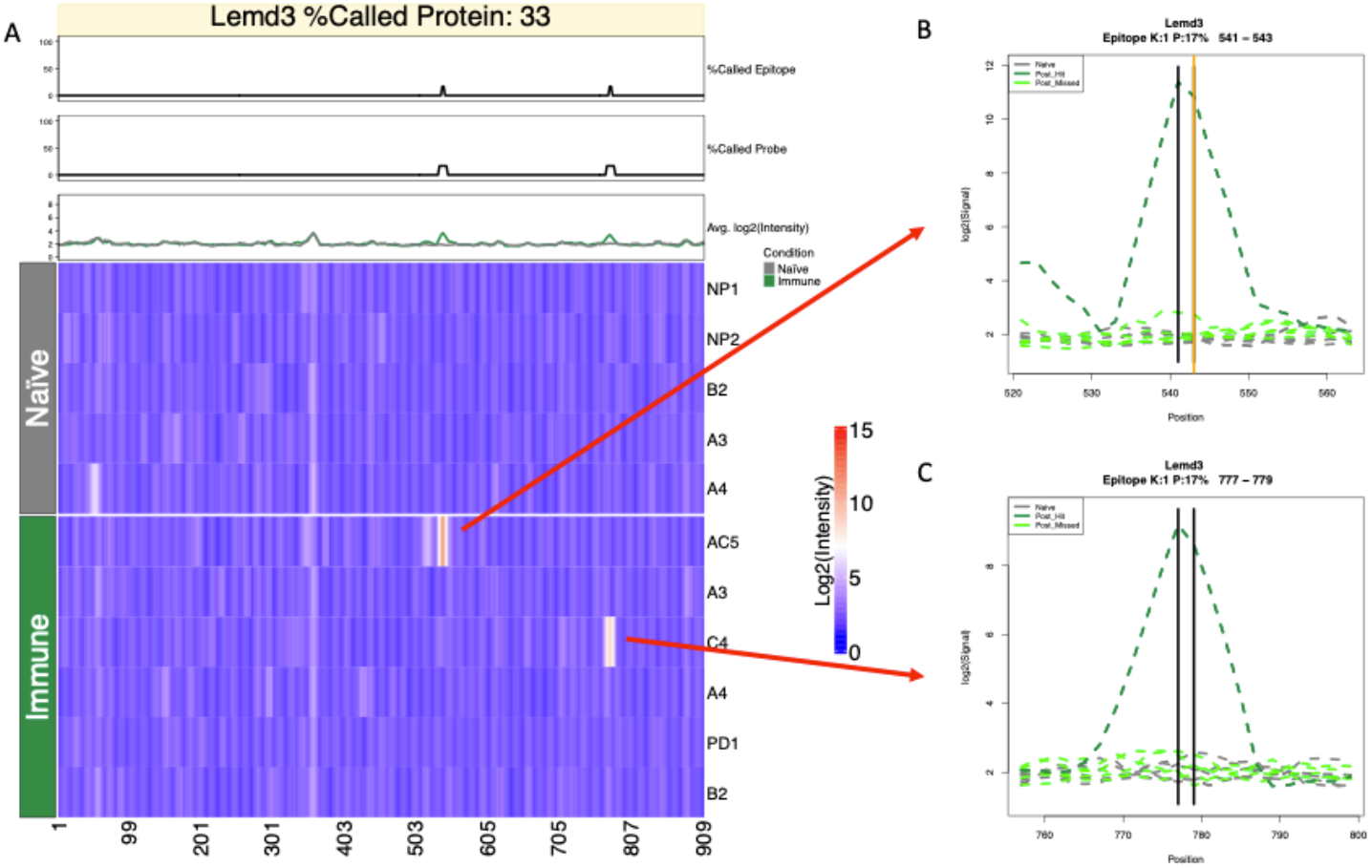
Heatmap and line plots of Lemd3. (A) Line charts and Heatmap of Lemd3 using normalized and smoothed intensity values. The X-axis indicates the starting position of the probe within the protein and the Y-axis rows consist of the individual pre-treatment (Naïve) and post-treatment (Immune) samples. The first line chart above the heatmap indicates the percent of positive samples that were called on the epitope-level (Moderate Significance), the second line chart indicates the percent of positive samples called at the probe-level (Moderate Significance), and the third line chart shows the average signal between negative (Naïve) and positive (Immune) samples. (B) Line plot of epitope detected for the mouse (AC5) sample. (C) Line plot of the epitope detected in the mouse (C4) sample. For both (B) and (C), the Y-axis is the normalized intensity values and the X-axis is the starting position of the probe within the protein and the dark green dotted lines indicate the samples that were called within the epitope boundary, which are indicated by vertical black or orange lines. The orange line indicates a probe tested and validated by ELISA.

### 3.4 Validation by ELISA from expert selected peptides

For the Melanoma dataset, the probe, epitope, and protein calls using the one-hit filter, unique segmentation method, the Wilkinson’s max for epitopes, and min+bonf for proteins with the inclusive, moderate, and restrictive significance levels were used to select peptides for validation. Hoefges et al 2023 selected 16 peptides (16-mers that had been tested in the high-density array) to validate using ELISA; 14 were selected based on their strong signal > 6SD over the mean by at least 3 of the 6 immune serum samples tested in the high-density array. The other 2 were selected as negative controls, based on signals < 3SD over the mean for all 6 immune serum samples in the high-density array. These were all then tested in a standard ELISA assay, with data reported out as optical density (O.D.) values for each serum sample (naïve or immune) tested, as detailed by Hoefges et al 2023 [29].

The replicates for ELISA were first averaged together. To indicate a positive hit, a threshold O.D. value of greater than or equal to two was used on each ELISA data point. For each peptide, the fraction of positive hits was calculated for the immune samples in the original and validated ELISA set and for the pre-/Naive samples in the validated set. A peptide was called validated for positive reactivity if 25% or more of the respective pre-/Naïve or post-/Immune samples were called.

The Internal Validation Cohort consisted of immune and naïve sera from the original 6 immune mice used in the original high-density array (and shown in Figure 5). When these sera were tested in the ELISA on these 16 peptides (**Figure 7A**), 10 out of 14 positively-selected peptides (71%) validated for positive reactivity on the immune samples, while 0 (0%) of the 14 peptides validated for positive reactivity on the naïve samples, as expected. In contrast, 0 out of 2 negatively-selected peptides (0%) had validated for positive reactivity on the immune samples, and similarly 0 out of 2 negatively-selected peptides (0%) had positive reactivity on the pre-/naive samples, as expected.

**Figure 7.**
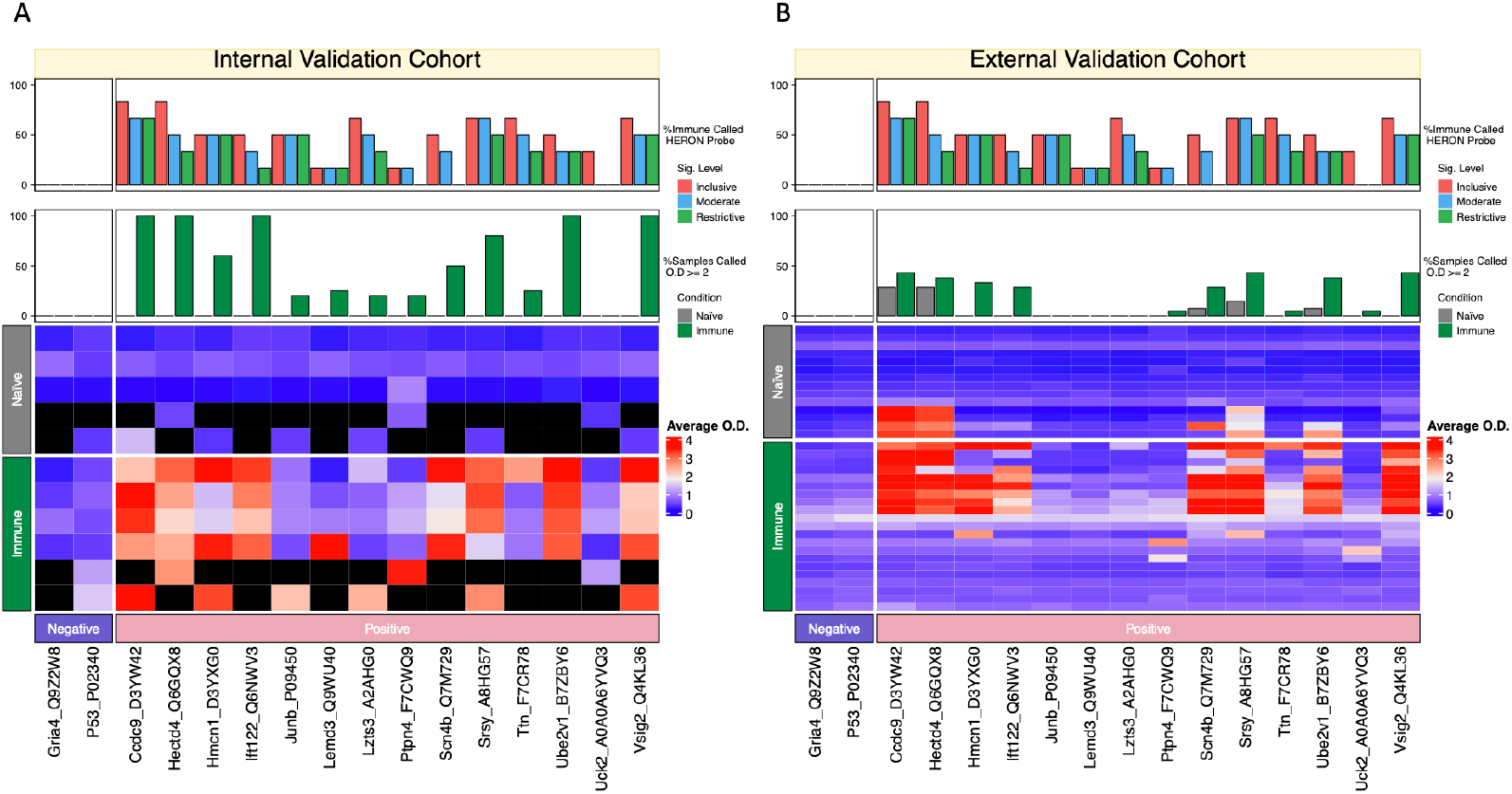
Heatmaps of ELISA results. Sixteen-mer peptides tested are shown on the columns and the individual naïve and immune serum samples are indicated on the rows. The top bar plots indicate the percentage of calls made on the probe-level by HERON in the Inclusive, Moderate, and Restrictive significance levels based on the original high density peptide array data (note these data are shown for comparison, and the graph of these high-density data are replicated for comparison’s sake in the top panels for A and B). The lower bar plot indicates the percentage of Naïve (grey) and Immune samples (green) that had an average ELISA value ≥ 2 O.D. intensity. Below these are the heat maps, showing the average across replicates O.D. values for each serum sample (naïve or immune) tested against each of the 16 peptides, with the 2 negative peptides shown at the far left and the 14 positive peptides at the right. (A) Internal Validation Cohort, (B) External Validation Cohort. Missing values (due to insufficient volume of serum available for that sample) are indicated as black values in the heatmaps.

The External Validation Cohort consisted of serum samples from 20 separate immune mice and naïve serum samples from 14 of those mice, that had not ever been tested before in any high-density array or any ELISA assay. When these 34 sera were tested in the ELISA on these 16 peptides (Figure 7B), most (8 out of 14) positively-selected peptides (57%) validated for positive reactivity on the immune samples, as expected, while few (only 2 of the 14) peptides (14%) had positive reactivity on the naïve samples, as expected. In contrast, 0 out of 2 negatively-selected peptides (0%) validated for positive reactivity on the immune samples, and similarly 0 out of 2 negatively-selected peptides (0%) validated for positive reactivity on the pre-/naive samples, as expected. These results with this external, independent, validation cohort are showing a similar, but not quite identical, pattern as seen with the internal validation cohort; the only difference is some of the naïve sera of these 14 independent cohort mice tested were also detecting a fraction of the positive peptides, which was not seen with naïve sera from the original cohort of 6 immune mice. The naïve sera from the original 6 mice did recognize some peptides in the high density array, but these peptides were not chosen for this validation ELISA testing. Thus, it is not surprising that some of the naïve sera in the independent cohort of mice might be able to recognize some peptides not recognized by the naive sera of the original cohort (**Figure 7**).

## 4. Discussion

Ultra-dense linear peptide binding arrays are useful for identifying the landscape of antibody binding to entire microbe or mouse proteomes for immune studies, however, many factors determine the interpretation of immune response in the form of antibody binding to linear peptides from peptide arrays. Defining antibody binding epitope boundaries can be challenging as regions of interest can be defined as regions with similar antibody binding responses across multiple samples, or a region from an individual or a small number of samples. We developed HERON, an algorithm and software package that can make antibody binding calls and estimate significance at the peptide, epitope, and protein level, while identifying global and individual features.

### 4.1 COVID-19 dataset performance

In the COVID-19 comparison between HERON and the method described in Heffron et al. 2021, we found good agreement between the probe- and protein-levels found by HERON. However, there were significant disagreements at the epitope-level. The discrepancies are due to differences in the way the epitopes are found and scored. The HERON method finds epitopes with a procedure that tries to balance an individual sample and across-sample levels and reports K of N or fraction scores to allow the user to determine the number of samples that are needed for valid epitopes. In contrast, the t-test method presented in the Heffron et al. 2021 study is looking for a difference of means between the COVID-negative and positive samples. Looking at the figures of epitope annotations for the spike protein of SARS-CoV-2 depicted in **Figure 3**, some of the epitope boundary differences between the Heffron et al. 2021 t-test method [11] and HERON method are probably due to some samples with a high post score. The t-test method makes calls based upon the average across all post samples, where a few outliers from the mean could result in a higher overall mean. The HERON method makes a call on each post sample, which could separate high binding “outliers” from lower binding “inliers”. The HERON method can also find probe hits similar to pepStat, which is a well-documented and tested software toolkit for analyzing peptide array data within the Bioconductor R environment. Most of the differences between pepStat and HERON can be attributed to edge effects, where a uncalled probe lies at the ends of epitopes that were previously found in Heffron et al. 2021. As mentioned previously pepStat is designed to call peptide sequences from multiple variants of the same protein, and uses a non-parametric approach to calculating FDR from the subtracted results of pre- and post-samples whereas HERON is a parametric approach using a differential t-test in the case of the COVID-19 and can handle whole proteome probe designs.

What parameters to use depends upon the type of question and goals of the study. In the case of virus epitope discovery, finding consistent epitopes across positive samples is desired to find a general biomarker for the development of vaccines and diagnostics. HERON also provides the ability to find positive sample specific epitopes and for finding epitopes of longer span using the “unique” segmentation method. Finding longer epitopes may be of interest to researchers when trying to find positive sample-specific epitopes.

### 4.2 Melanoma dataset performance

Our results using the different meta p-values on the Melanoma dataset gives an idea of what method to use when calculating an aggregate p-value for the epitope and protein level calls. Different methods need to be employed depending upon whether the desired result is for all p-values within the aggregate to be significant (epitope level) or at least one of the p-values within the aggregate need to be significant (protein level). We have chosen a few different meta p-value methods to compare and contrast against each other. Using Wilkinson’s max p-value is also competitive for epitopes. In the literature, there are many more meta p-value methods and a comparison of those methods with the ones studied here would be a good next step. However, some of them, while modeling the dependencies between the p-values using correlation coefficient, require many more samples to be accurate [22, 42] Berk-Jones, HC, and others take a longer time to run [21].

From our results, the fisher method seems competitive and the extensions to that method such as the empirical Brown [22] methods that use covariance between samples to estimate p-values might be helpful in finding consistent probes, epitopes, or proteins across all samples when the number of samples is high. As the antibody repertoire of mice shows stochastic variability between mice, based on VDJ recombination of genes determining antigen binding immunoglobulin regions during immune ontogeny, the number of probes, epitopes and proteins expected to be recognized by serum from all immune mice is expected to be small. Our initial results evaluating this, using the methods developed herein, confirms this prediction [29]. However, we are finding that there are some probes, epitopes and proteins co-recognized by a substantial fraction of immune mice, suggesting some antigens may be of importance in a substantial fraction of mice. For epitope meta p-values, we will use methods that ensure that the underlying p-values are mostly significant for epitope scoring. The one-hit filter seems to increase the correlation results slightly at the cost of reduced number of hits found. Although the hits that were missed may be due to noise from the peptide array. The choice of segmentation algorithm depends upon the desired type of epitopes. The “unique” segmentation method seems to find on average longer runs of probes and epitopes, while the other clustering methods seem to find shorter and more consistent epitopes. In the case of the Melanoma dataset, longer runs of peptides for epitopes seem to be desired, whereas the COVID-19 and other virus type studies trying to find good vaccine targets would prefer the shorter and more consistent sequences.

### 4.3 Assumptions, Issues, and Calibration

There are several complexities with running HERON during processing peptide binding array datasets and we list the majority of them here. Starting at the unique sequence-level, HERON assumes that the data is normally distributed, or if the data has been smoothed, the smoothed probe-level is normally distributed.

#### 4.3.1 Assumptions on the unique sequence or smoothed probe-level

For calculating the differential t-test p-values/scores, HERON assumes that the variance of the post-sample is the same as the estimated variance of the pre-sample and that the post-sample signal represents the mean of the post-group values. To help deal with the higher variance and the obvious breakdown of the assumptions, we use the degrees of freedom from the pre-samples to estimate the one-sided p-value from the t-distribution.

Upon calculating the global z-test p-values/scores, HERON assumes that the sequence or smoothed probes signals are comparable across different sequences and that the mean and standard deviations can be used to calculate the p-value against one sequence signal for a post sample.

When combining the differential and global z-test p-values, the Wilkinson’s max meta p-value method assumes independence among the p-values to be combined. More study is needed to determine if HERON has violated this assumption and, if so, the ramifications when estimating the combined p-values/scores or if using a different meta p-value that more tolerant to correlated p-values would improve HERON’s estimation of significance on the probe-level.

Finally, in the case of copying the unique sequences to the probe-level, there is an assumption that it is fair to do so. The main assumption is that resulting probe-level p-values are still well behaved. In the case of the Melanoma dataset, since we are smoothing the data beforehand, each probe can be treated as an independent measurement, even though there are some peptide probes that come from a sequence that maps to more than one protein. The COVID-19 dataset, however, is unsmoothed and contains many sequences shared between proteins from different strains of the coronaviruses. To process unsmoothed data in HERON, we estimate the adjusted p-values on the unique sequences and then copy the result to the respective probe-level identifiers.

#### 4.3.2 Assumptions on the epitope-level

For the epitopes, or group of consecutive probes across a protein, the meta p-value methods used assume that the p-values are accurate and well-calibrated. Several of them (Wilkinson, min+bonf, Fisher, etc.) assume independence among the p-values, which is directly violated as we group adjacent probes which overlap, as epitopes. Other methods such as the Cauchy combination test or the harmonic mean try to alleviate this assumption, while other meta p-value methods (Brown, Kosts, etc.) attempt to model the covariance between the p-values to improve the estimation.

There is also an assumption that the epitopes identified by the epitope finding methods are the only ones to be included in the analyses. For example, do we need to correct for the number of all possible epitopes per protein? Currently, HERON treats the list of epitopes as a separate list and just corrects using the Bonferroni-Hochberg algorithm when reporting the adjusted p-values for the epitopes and uses the uncorrected p-values for estimating the protein-level p-values.

#### 4.3.3 Assumptions on the protein-level

Going up another level in the hierarchy of probes, epitopes, and proteins, the protein p-values calculations assume that the epitope regions and p-values/scores are well-calibrated and well-behaved and the further assumptions made by the meta p-value method used to estimate the protein p-value/scores from the epitope p-values. We also assume that the adjusted p-value correction only needs to be applied to the proteins for which at least one epitope region was found.

#### 4.3.4 Calibration of p-values using permutation tests

One of the ways to alleviate some of the issues presented above is to re-calibrate the scores into actual accurate/well-behaved p-values. Permutation statistics, shuffling the sample labels, calculating the resulting p-values/score, and using the permutation scores to calibrate the raw p-values is a popular method for ensuring well behaved statistics.

While the use of permutation statistics to provide accurate p-values on the unique sequence or probe-level is seemingly straightforward (Data not Shown), it is unclear how to properly perform this calibration at the epitope, and protein levels. We are currently exploring this avenue to see if providing calibrated p-values to the unique/smoothed probe-level also equivalently gives well calibrated scores at the epitope and protein level, or if more complicated calibration is needed.

Furthermore, permutation statistics is also affected by the number of samples used in the experiment. For example, in the Melanoma experiment, there are 8,459,970 unique peptide probes (not 6,090,593 due to the smoothing across probes), and only 11 biological samples. Testing for all possible permutations (11! = 39,916,800) would achieve a minimum permutation estimated p-value of 2.505×10^−8^, which after using a Bonferroni correction against all probes would achieve a corrected p-value (2.505×10^−8^ x 8,459,970) of 0.212 or 0.153 using 6,090,593 unique sequence probes. While there are methods to further increase the p-value accuracy in the lower range of p-values [43] with fewer permutations, there is a limitation with using permutation tests with millions of features and a small set of independent biological samples.

Finally, recent studies have shown that care must be taken when using the permutation test techniques [44]. In this paper, we use the normalized log-transformed data, which is part of the suggested use of the Box-Cox transformation [45]. Other methods for analyzing high-throughput peptide binding array data perform a log-log transformation before estimating the statistics [13, 15]. While studying the application of permutation calibration, we will investigate the usefulness of the Box-Cox transformations when calculating the statistics and calibration procedures.

### 4.4 Proposed Extensions

With our initial success of the workflow for finding probes, epitopes, and proteins in a high-throughput protein data set, several avenues exist to further improve upon the HERON method. In the next few paragraphs, we discuss four ideas: 1) Improving the combination of probe-level p-values with copulas or trying other meta p-value methods for calculating epitope and protein p-values, 2) Enhancing the flexibility of K of N calls using a Binomial or Poisson Binomial, 3) Trying different segmentation approaches for finding epitopes such as bi-clustering, and 4) using Bayesian models to model the hierarchical nature of the calls made by the workflow.

HERON allows for the combination of probe-level p-values from the global and differential tests by using Wilkinson’s max meta p-value method, which may have violated the independent p-value assumption. Other meta p-value methods exist that are tolerant to dependent p-values [19, 23] and could be used to estimate the combined p-value. Other meta p-value methods incorporate correlation to model the dependence [22, 46], however, these methods’ accuracy is determined by the number of samples for estimating the covariances [36].

As previously mentioned, HERON was not designed to handle cases with a high incidence where one amino acid sequence maps to many proteins and further study is needed to investigate how these redundancies affect the scores and how to adapt HERON to improve handling of datasets with this feature.

The current workflow calculates the K of N for each probe, epitope, and protein after the respective FDR cutoff is chosen. A user may want to infer a statistical test for selecting elements where K > b. Using the Poisson binomial distribution, we could estimate a probability for each level of K to calculate p-values with a null hypothesis of K ≤ b. While another approach would be to just use a binomial with a set probability. More investigation is needed to determine the pros and cons of utilizing these statistical tests within HERON.

The segmentation methods tested could be extended with other clustering methods. Use of different distance calculation functions, clustering score methods or clustering methods could further improve discovery. The hierarchical and skater clustering mentioned here were only run using average Hamming or Euclidean distance. There are many more cluster distance metrics available (e.g., Manhattan) to compare and contrast. In this paper, HERON finds the optimal clustering by finding the number of clusters/cuts by optimizing the average silhouette score. Other clustering scoring metrics, such as the gap statistic [47, 48] could also be used. The problem with the gap statistic is the increase in processing time to run the bootstraps for finding the optimal clustering. Using bi-clustering with the skater graph-like constraints would allow segments to be found that could overlap with other segments. Future iterations of HERON may include some of these and more study would be done to determine the advantages and disadvantages of using these different clustering approaches.

The provided workflow attempts to place significance values on epitopes, proteins, and probes using meta p-value methods. Another approach would be to estimate probabilities at each level jointly using all the provided data.

Implementing a Bayesian hierarchical model, which can borrow information from the estimated probability of the different levels from a fitted Bayesian model, would be a natural extension of the workflow presented in this paper. For peptide antibody array analysis, Bayesian models have been proposed. The pepBayes implementation is designed to handle one protein at a time from multiple strains and does not take into account the sequential probes across the protein [3]. Another Bayesian model does include provision of handling sequential probe signals by using a latent autoregressive component, however the authors propose a limit in the problem size (300 peptides and 50 samples) when using their implementation in WinBUGS [17]. Additionally, neither consider cases for multiple proteins and the additional information that can be obtained and leveraged for estimating parameters on the protein level.

## 5 Conclusions

Ultra-dense peptide binding arrays are powerful tools for studying the abundance of different antibody repertoire in serum samples to understand adaptive immune responses. HERON, an R package for analyzing peptide binding array data, is a flexible and powerful tool for selecting groups of linear peptide probes with improved reliability and reproducibility when considering epitopes rather than single peptide probes due to several factors; first, there are many more probes than epitopes in the proteome, giving a larger number of possible mismatches. Second, an individual epitope can be a component of several overlapping probes; our HERON algorithm for detecting epitopes recognized by separate assessments of serum samples, requires a degree of similar recognition of the related epitope containing probes by the 2 samples, but does not require complete identity of probe recognition and signal. This enables higher reproducibility of epitopes recognized with high signals than peptides recognized with high signals when replicate chips are evaluated for separate aliquots of the same immune serum sample when evaluating proteins that are recognized, since a single protein might be recognized by different individuals at different regions.

With HERON, finding interesting linear peptide probes, epitopes, and proteins within experimental data from large (mouse proteome) and small (COVID-19 proteins) is possible. We have plans to incorporate HERON in the Bioconductor environment and to maintain it to be of use to the scientific community. Future improvements to HERON are proposed and other approaches will be explored to provide versatile algorithms for analyzing peptide binding arrays in a multitude of possible experiments.

## 6 List of Abbreviations

BH: Benjamini-Hochberg
HERON: **H**ierarchical antibody binding **E**pitopes and p**RO**teins from li**N**ear peptides
O.D.: Optical density
FDR: False Discovery Rate
a.a.: amino acid
Ab: Antibody

## List of Figures and Tables

**Supplementary Figure 1 – Probe estimated p-values vs normalized signal for one representative serum sample from an immune mouse (AC5) included in the melanoma dataset using moderate statistics parameters**.

**Supplementary Figure 2 – Technical replicate scatterplots of -log**_**10**_ **of adjusted p-values on probes, epitopes, and proteins using the moderate level statistics**.

**Supplementary Figure 3 – Heatmap and Lineplots of Hmcn1**.

**Supplementary Table 1 - Comparison of Probe Calls between the HERON, pepStat, and Heffron et al. 2021 Methods**

**Supplementary Table 2 – Comparison of Epitope Calls between HERON and Previous Method**.

**Supplementary Table 3 – Technical Replicate Correlation Results using Different Parameters for HERON**.

**Supplementary Table 4 – Technical Replicate Correlation Results on the Probe-level**.

**Supplementary Table 5 – Technical Replicate Correlation Results Averaging across Same and Different Arrays and then Statistical Parameters**.

## Acknowledgements

We would like to thank Ken Lo, John Tan, Jigar Patel, Trang Le and Jess Vera for helpful conversations and assistance in testing the HERON package.

## Author Contributions

Conceptualization: SJM, AH, PS, IMO. Data curation: SJM, AH. Formal analysis: SJM, AH. Funding acquisition: PS, IMO. Investigation: SJM, AH, AKE, KL, PS, IMO. Methodology: SJM, AH, AKE, PS, IMO. Project administration: SJM, IMO. Resources: PS, IMO. Software: SJM, IMO. Supervision: SJM, PS, IMO. Validation: SJM, AH, IMO. Visualization: SJM, PS, IMO. Writing – original draft: SJM, IMO. Writing – review & editing: SJM, AH, AKE, KL, PS, IMO.

