## Supplementary Figures for "Ranking Antibody Binding Epitopes and Proteins Across Samples from Whole Proteome Tiled Linear Peptides"

[1]Department of Biostatistics and Medical Informatics, University of Wisconsin-Madison, WI, [2]University of Wisconsin Carbone Comprehensive Cancer Center, University of Wisconsin-Madison, WI [3]Department of Human Oncology, University of Wisconsin-Madison, WI, [4]Department of Pediatrics, University of Wisconsin-Madison, WI, [6]Department of Obstetrics and Gynecology, University of Wisconsin-Madison, WI, [6]Center for Human Genomics and Precision Medicine, University of Wisconsin-Madison, WI

*To whom correspondence should be addressed.

### Supplementary Figures

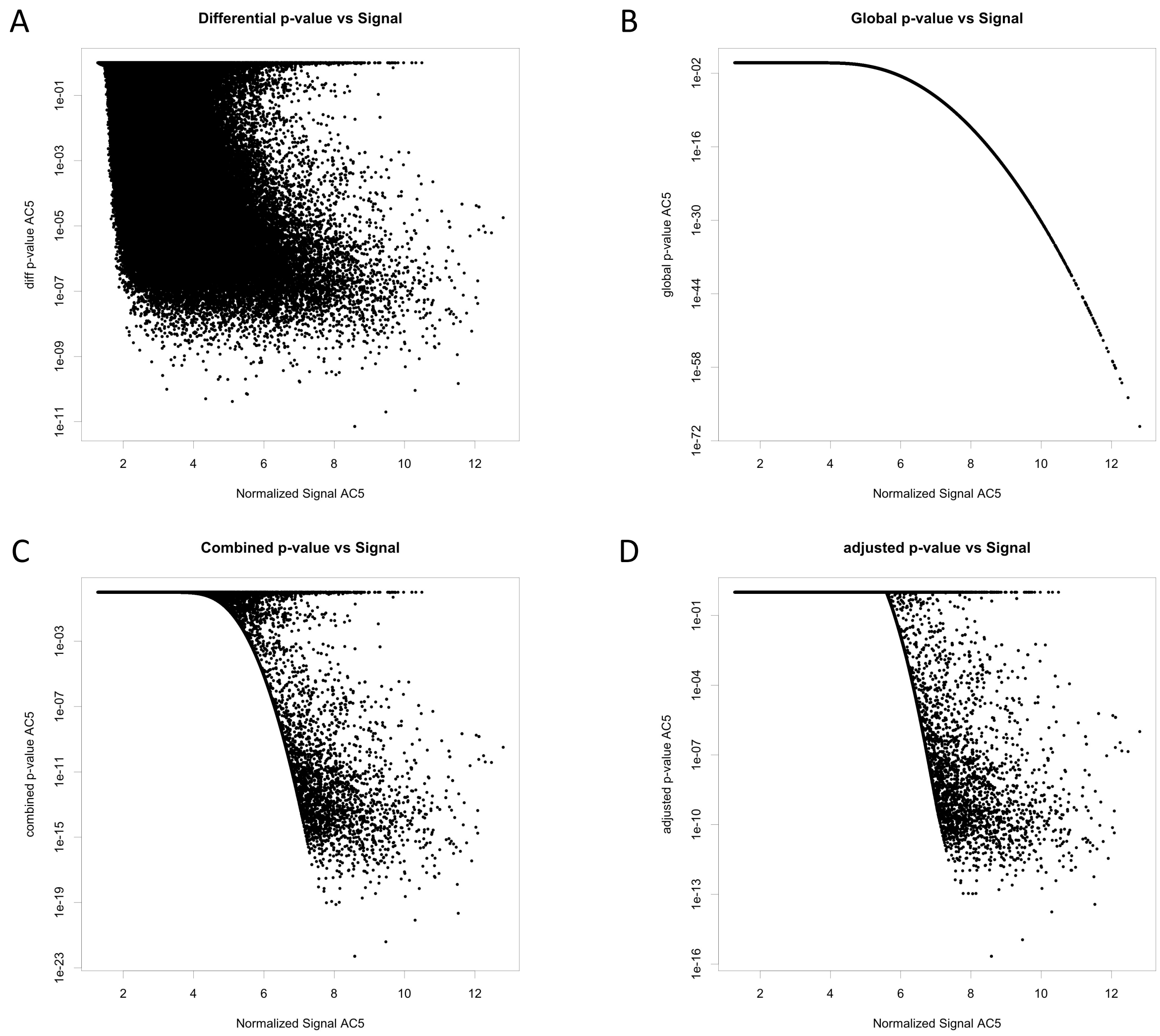


**Supplementary Figure 1 – Probe estimated p-values vs normalized signal for one representative serum sample from an immune mouse (AC5) included in the melanoma dataset using moderate statistics parameters.** In each panel, each dot represents the value obtained for this single serum sample on the ~$8 \times 10^6$ peptide probes (16-amino acids each) tested in the Nimble peptide array system. A) result of the differential p-value estimation using t-test. B) result of global p-value using z-test. C) result of the combined p-value using Wilkinson's max. D) result of adjusted p-values (Benjamini-Hochberg).

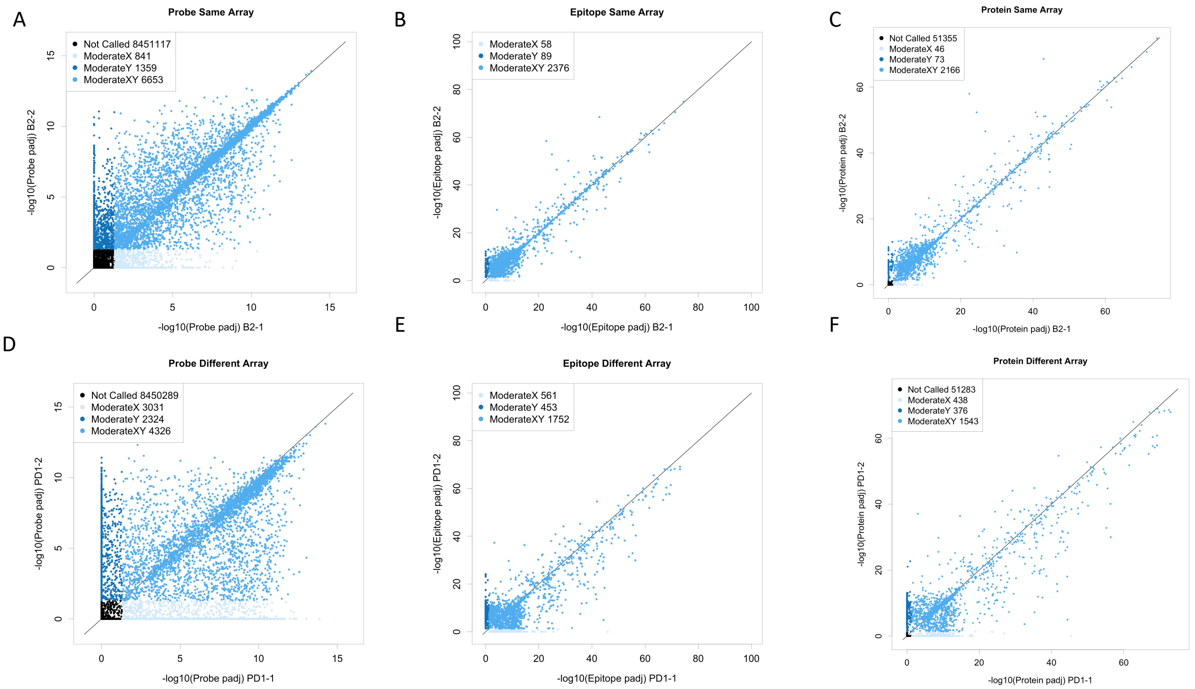


**Supplementary Figure 2 – Technical replicate scatterplots of $-\log_{10}$ of adjusted p-values on probes, epitopes, and proteins using the moderate level statistics.** A, B, and C are from the technical replicates collected on the same array (two separate replicate data sets obtained in the same assay using split serum aliquots from immune mouse B2). D, E, and F are from the technical replicates collected on arrays collected at different times (two separate replicate data sets obtained on replicate high density array chips in assays performed one-year apart, using split serum aliquots from immune mouse PD1). A and D are from the probe-level p-values, B and E are from the epitope-level p-values, and C and F are from the protein-level p-values. Light-blue are the probes, epitopes, or proteins that are called in just the 1st replicate shown on the X-axis (B2-1 or PD-1). Dark blue are the probes, epitopes, or proteins that are called in the 2nd replicate (B2-2 or PD-2) shown on the Y-axis. Medium blue are the probes, epitope, or probes that are called in both replicates (and thus running more on the diagonal). For A and D, the black points indicate probes that were not called in either replicate.

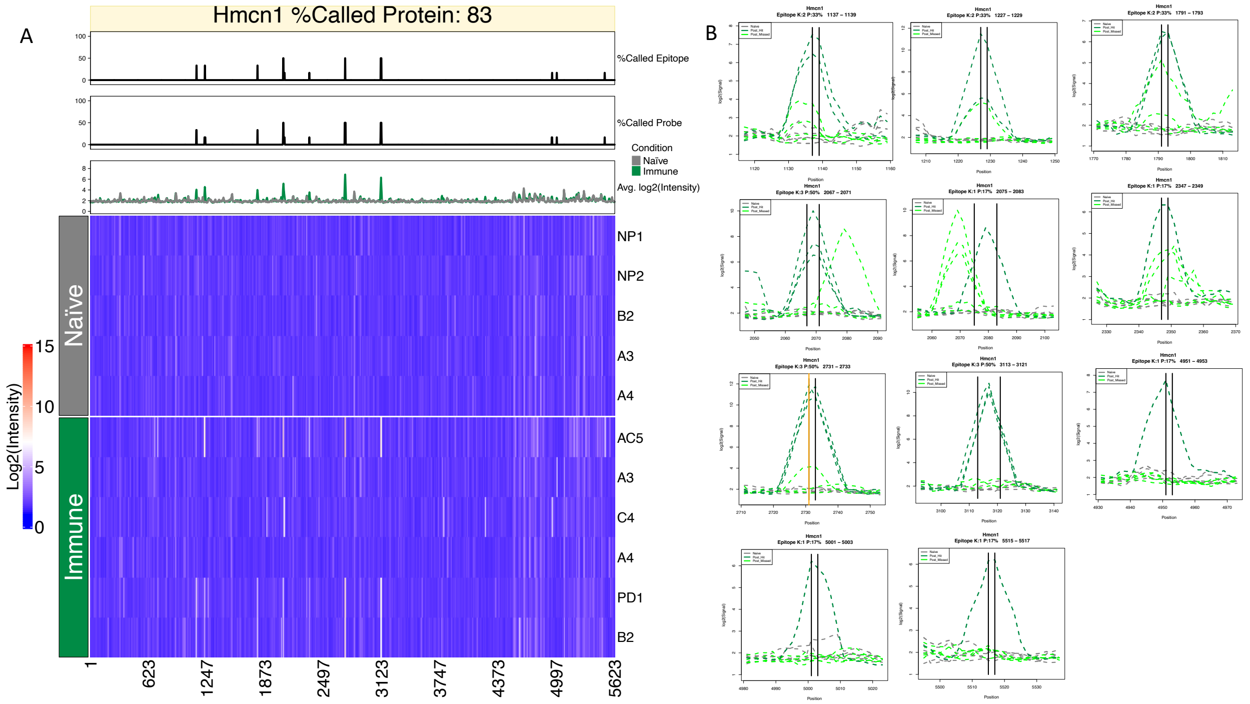


**Supplementary Figure 3 – Heatmap and Lineplots of Hmcn1.** (A) Line charts and Heatmap of Lemd3 using normalized and smoothed intensity values. The X-axis of indicates the starting position of the probe within the protein and the Y-axis is the individual pre- and post- samples. The first line chart above the heatmap indicates the percent of positive samples that were called on the epitope-level (Moderate Significance), the second line chart indicates the percent of positive samples called at the probe-level (Moderate Significance), and the third line chart shows the average signal between negative (Naïve) and positive (Immune) samples. (B) Line plots of 11 epitopes detected for the various mouse samples, Y-axis is the normalized intensity values and the X-axis is the starting position of the probe within the protein. The dark green dotted lines indicate the samples that were called within the epitope boundary, which are indicated by vertical black or orange lines. The orange line indicates a probe tested and validated by ELISA.
